# Loss of *Ag85A* disrupts plasma membrane domains and promotes free mycolic acid accumulation in mycobacteria

**DOI:** 10.1101/2025.08.05.668640

**Authors:** Takehiro Kado, Shivangi, Jake Jordan, Joel S. Freundlich, M. Sloan Siegrist, Yasu S. Morita

## Abstract

The mycomembrane of mycobacteria, composed primarily of long-chain mycolic acids, is critical for cell survival, structural integrity, and resistance to environmental stress, yet its underlying synthesis mechanisms remain incompletely understood. This study investigates the role of Ag85A, a key enzyme in mycomembrane synthesis, in regulating plasma membrane domains and cell envelope organization in *Mycobacterium smegmatis*. Using Δ*Ag85A* deletion mutants, we combined microscopy, biochemical assays, thin-layer chromatography, and lipid analysis to evaluate changes in membrane structure, chemical accumulation, and lipid composition. *Ag85A* deletion leads to altered plasma membrane domain organization, increased chemical accumulation, changes in cell envelope lipid composition. Unexpectedly, lipid analysis revealed accumulation—not depletion—of mycolic acids in the mutant, suggesting that increased permeability is not directly due to mycolic acid loss. These findings highlight a novel link between mycomembrane composition and plasma membrane domain stability. Our study not only advances understanding of mycobacterial cell envelope architecture but also identifies potential targets for enhancing drug penetration in resistant mycobacterial infections.

**Significance:** The unique cell envelope of mycobacteria is central to survival, enabling it to resist immune defenses and antibiotic treatment. In this study, we reveal a novel function of Ag85A, a synthase of outmost layer of mycobacteria beyond its known role in mycolyltransferase activity: it is essential for the formation of plasma membrane domains that orchestrate cell envelope synthesis. By structuring plasma membrane domains, Ag85A contributes directly to the resilience of the cell envelope, reinforcing mycobacterial survival mechanisms under hostile conditions. Our findings provide a pivotal insight into mycobacterial cell biology, with broad implications for therapeutic development. Disrupting Ag85A-dependent membrane domain formation could weaken the protective cell envelope, offering a promising approach to enhance current treatments.

## Introduction

The outer membrane in diderm bacteria provides structural integrity and stability to the bacterial shape (1), acts as a barrier against antibiotics (2), and elicits immune responses while inhibiting lysosome fusion to protect bacteria from host defenses (3). In mycobacteria, this outer membrane, known as the mycomembrane, is characterized by long carbon chain fatty acids called mycolic acids (4). Two key mycolic acid-containing molecules, trehalose mono-mycolate (TMM) and trehalose di-mycolate (TDM), play essential roles role in bacterial survival and pathogenicity (4–10).

TDM and TMM are crucial for maintaining the integrity and function of the mycomembrane as numerous important antibiotics, including isoniazid, SQ109, delamanid, PA-824, and ethambutol, are designed to interfere at different points in the synthesis pathway of TMM and TDM (11–15). Central to this process are the enzymes from the antigen 85 family, specifically Ag85A, -B, and -C. These enzymes are highly abundant, making up to 41% of the protein in culture supernatants, and they play a pivotal role in the synthesis of TMM and TDM, as well as the attachment of mycolates to arabinogalactan (16, 17). Their abundant presence and critical function in mycolate metabolism make them attractive targets for understanding the role of mycomembrane in mycobacterial survival strategy. Among the isoforms (Ag85 A, B, C), Ag85A receives the most attention due to its highest transferase activity (18).

Research involving the knockout of *Ag85A* in *M. smegmatis*, *M. bovis*, and *M. tuberculosis* has revealed several phenotypic changes. Antibiotic sensitivity and accumulation increases in *M. smegmatis* and *M. tuberculosis* (19–21). Growth rates in macrophages is reduced in *M. bovis* and *M. tuberculosis* (22, 23), and biofilm formation is disrupted in *M. smegmatis* and *M. tuberculosis* (20, 24, 25). Additionally, transposon screening showed that deletion of *Ag85A* may delocalize plasma membrane domains (26), known as the inner membrane domain (IMD) (26–28) where cell envelope precursors are synthesized in *M. smegmatis* (27, 29). The previous research suggested that the membrane domain synthesize cell envelope including cell wall and domain in mycobacteria, diderm bacteria and plant cells (30, 31). Those facts suggest that Ag85A, in turn outer membrane, is able to feedback plasma membrane as well to form the plasma membrane domain.

This study elucidated the essential role of the mycomembrane layer in mycobacterial survival, with a focus on characterizing the deletion mutant of the mycomembrane synthase Ag85A to gain deeper insights into mycomembrane function. Ag85A, along with other key genes such as *ponA2* (a cell wall synthase) and *cfa* (a C19-methylated fatty acid synthase), plays a crucial role in plasma membrane domain formation in *M. smegmatis*. As the deletion mutants of *ponA2*, *cfa*, and *Ag85A* exhibit significant disruptions in biofilm formation, impaired growth in macrophages *in vitro* and in mice *in vivo*, and increased drug sensitivity (20, 22–25, 32–34), these findings underscore the critical role of plasma membrane domains in maintaining cellular homeostasis and survival, and they highlight these domains as promising targets for new drug development.

## Result

### Accumulation of membrane fluidizer in ΔAg85A

Since transposon sequencing identified *Ag85A* in the dibucaine, the deletion of Ag85 could sensitize *Mycobacterium smegmatis* to dibucaine or the deletion delays the establishment of the membrane domain during the recovery incubation. To determine if Δ*Ag85A* is sensitized to the dibucaine treatment or delayed from the recovery, the deletion mutant was treated with dibucaine for three hours and the colony forming unit was assessed after the treatment. Δ*Ag85A* showed the smaller number of colony-forming unit (CFU) after the treatment (Fig1A). The deletion of *Ag85A* sensitized *M. smegmatis* to dibucaine.

While the deletion increases the dibucaine sensitivity, we still do not know what the cause of death in the mutant. Since Ag85A is one of the main enzymes developing outer mycomembrane of mycobacteria, the absence of Ag85A can increase mycomembrane permeability resulting in the increased dibucaine accumulation in the cell. We quantified dibucaine in the cell using high-performance liquid chromatography mass spectrometry (HPLC-MS). We treated wild-type and Δ*Ag85A* with dibucaine, and the dibucaine taken up by the cells were extracted by a mixture of methanol, acetonitrile, and water (MAW) (2:2:1, vol/vol/vol). HPLC-MS analysis revealed significant accumulation of dibucaine in the Δ*Ag85A* mutant (Fig. 1B). However, no significant difference was observed between the deletion strain and the complemented strain (Fig. 1B). Potential matrix effects among different matrices of strains were evaluated but found to be negligible (Supplementary figure 1). The incomplete recovery of drug accumulation in the complemented strain may reflect partial restoration of colony-forming ability (Fig. 1A); while the number of CFUs increased, the colony size remained smaller than that of the wild-type.

**Figure 1.**
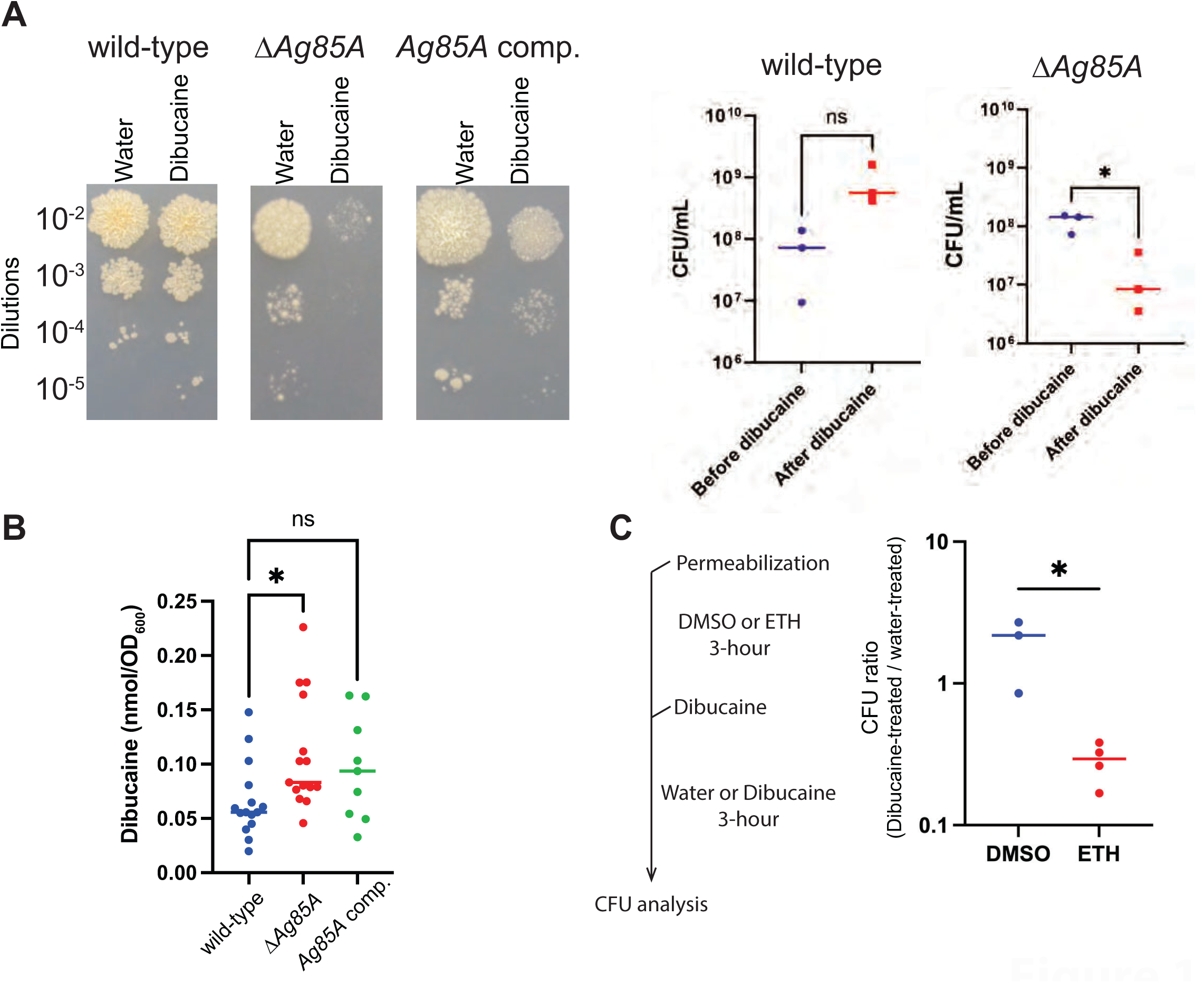
Ag85A protects *M. smegmatis* from the membrane fluidizer accumulation. (A) Left, the survival of Δ*Ag85A* after 3-hour treatment of water or dibucaine treatment was analyzed on the 7H10 plate. Right, colony-forming units (CFU) were calculated from three biological triplicates. There were no statistically significant differences in wild-type (indicated as ns) but Δ*Ag85A* showed the statistical differences (* P<0.05) as determined by Mann-Whitney U test. (B) Dibucaine accumulation in cell envelope was quantified by liquid chromatography-mass spectrometry. p-values were determined by Kruskal Kruskal-Wallis test with Dunn’s multiple comparisons test from three technical replicates from three biological replicates. replicates ns p= 0.4936, p=0.0323 (C) Cells were pretreated with either ethambutol or DMSO (vehicle control) for 3 hours. After the pre-incubation, the cells were washed then treated with dibucaine or water for an additional 3 hours. CFU counts were obtained for each group both before and after dibucaine treatment, and CFU ratios were calculated by dividing the CFU of the dibucaine-treated group by the CFU of the water-treated group. P=values were determined by Mann-Whitney U test. * p=0.0470.

Dibucaine accumulation may result from increased mycomembrane permeability caused by the deletion of *Ag85A* (21). This observation led us to hypothesize that alternative methods of permeabilizing the mycomembrane could similarly promote dibucaine accumulation within the cell, ultimately resulting in bacterial death. To assess whether pre-permeabilization of the mycomembrane increases the susceptibility of *M. smegmatis* to dibucaine, we conducted a two-step experiment. Cells were first treated with either ethambutol, a known molecule inhibiting mycomembrane biosynthesis (4, 21, 35–39), or DMSO as a control, followed by dibucaine or water treatment. Susceptibility was assessed by calculating the ratio of CFU in the dibucaine-treated group relative to the water-treated group. A CFU ratio less than 1 indicates that pre-permeabilization reduced cell viability in the presence of dibucaine. Treatment with DMSO showed no significant difference in CFU counts between dibucaine and water treatments, suggesting no sensitization under these conditions. In contrast, pretreatment with ethambutol resulted in a marked decrease in CFUs upon dibucaine exposure, yielding a CFU ratio below 1 (Fig 1C). These results indicate that ethambutol-mediated mycomembrane permeabilization sensitized *M. smegmatis* to dibucaine, enhancing its antibacterial effect.

### Delocalized membrane domain under knockout and knockdown of Ag85A

High dibucaine sensitivity and accumulation in Δ*Ag85A* suggested that the previous Tn-seq identified *Ag85A* because the permeabilized mycomembrane allowed the accumulation of excess amount of dibucaine in the cell but the relationship between *Ag85A* and the localization of membrane domain remains unknown. To observe membrane domain formation, *Ag85A* depletion strain was generated in the background of *M. smegmatis* strain (27) which expresses domain marker proteins: GlfT2 tagged with HA and mCherry, and Ppm1 tagged with cMyc and mNeonGreen. Overnight treatment with anhydrotetracycline was used for depleting *Ag85A*. Microscopic analysis after the depletion suggested that the sub-polar localization of the membrane domain was de-localized as the fluorescence protein localization was distributed along entire cell (Fig. 2A). The signal was quantified by Oufti and the signal peak was also lower in the depletion strain. To confirm whether this delocalization was caused by anhydrotetracycline itself, we added anhydrotetracycline to the parental strain—prior to insertion of the DNA region enabling *Ag85A* depletion—and observed no delocalization (Supplementary Fig. 2). Those data suggest that *Ag85A* is needed for the subpolar localization of plasma membrane domain (Fig. 2A). To further investigate domain formation, we performed biochemical analysis of the membrane domains in Δ*Ag85A* after lysing the cells through nitrogen cavitation, followed by sucrose gradient fractionation. IMD and PM-CW domains were fractionated in fractions 4-6 and 8-12, consistent with previous literature in the absence of dibucaine (Fig. 2B). However, following dibucaine treatment, the IMD domain was absent, and PM-CW appeared in fragmented distributions across fractions 2-3, 6-7, and 10-12 (Fig. 2B)—a pattern not observed in the wild-type (26–30). These findings suggest that the lateral separation of the plasma membrane in Δ*Ag85A* is more fragile and susceptible to disruption by membrane fluidizers compared to the wild-type.

**Figure 2.**
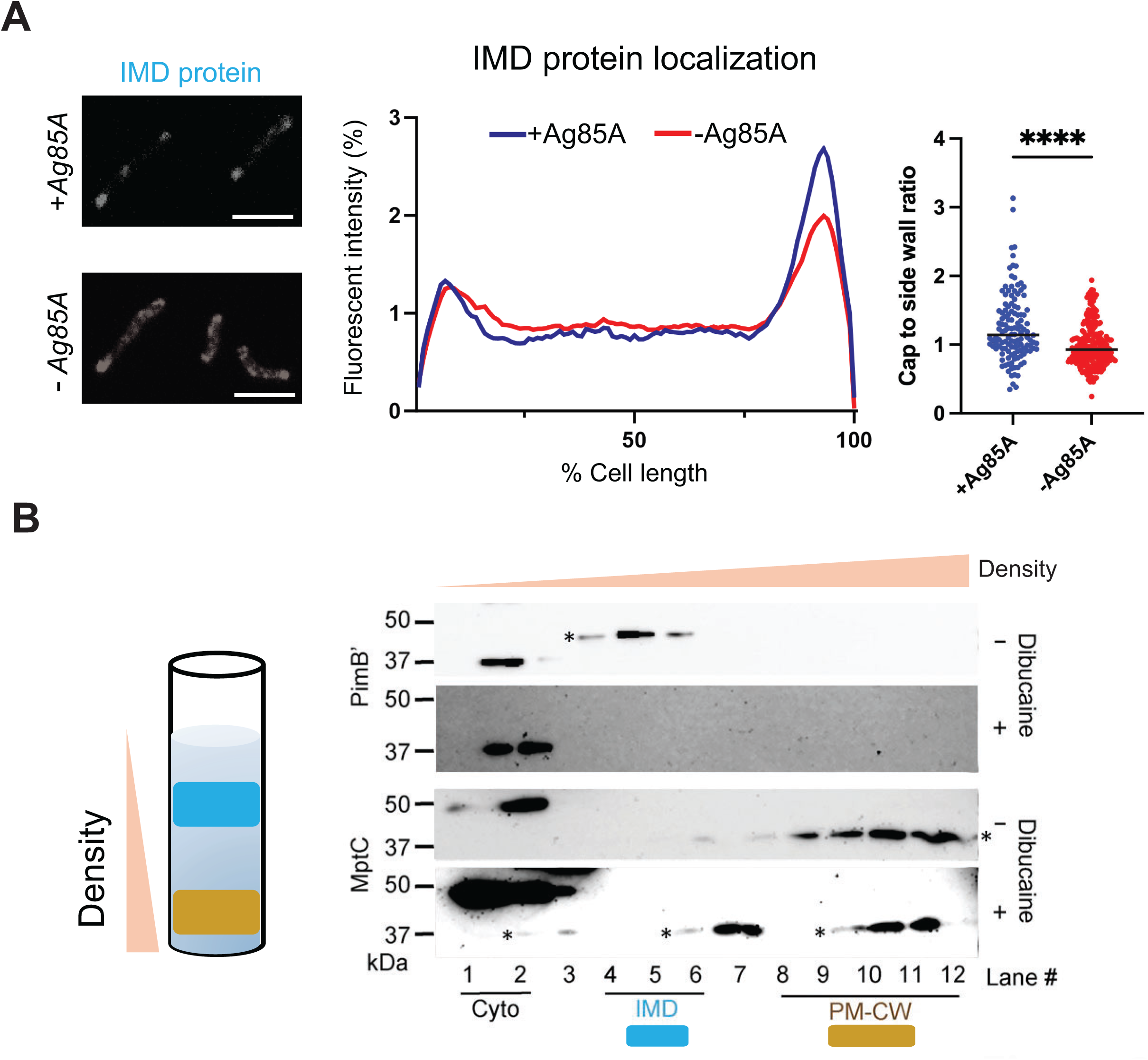
The depletion of *Ag85A* induced IMD delocalization. (A) Delocalization of mCherry-GIfT2 by the depletion of *Ag85A* in *M. smegmatis*. mCherry-GIfT2 was imaged after 12-hour anhydrotetracycline treatment. Images are representative of three independent experiments, and fluorescence distributions of the fusion proteins after chemical treatment were calculated from three independent experiments. Lines show the average of all cells (50 < n < 75). Signal was normalized to cell length and total fluorescence intensity. Scale bar, 5 μm. (B, left) Diagram of IMD and PM-CW fractionation after ultracentrifugation. (B, right) To visualize IMD and PM-CW, PimB’ and MptC which are established marker protein were analyzed by western blotting. Asterisks indicate the proteins analyzed.

### The accumulation of free mycolic acid led by the deletion of Ag85A in M. smegmatis

Cell wall and tuberculostearic acid in plasma membrane are important for establishing plasma membrane domain (29–31). As *Ag85A* enzymatically mediates the synthesis of TDM from TMM, or the reverse reaction (17), the deletion may affect mycomembrane composition and the disrupted structural integrity can be a leading factor of dibucaine accumulation and the delocalization of membrane domain. However, structural change of mycomembrane in *Ag85A* deletion is not completely understood. We further characterize mycomembrane biochemically to investigate how the deletion of *Ag85A* is affecting mycomembrane structure.

Whether the deletion of *Ag85A* compromises the mycomembrane remains debated. Nguyen et al. demonstrated higher release of TDM from the mycomembrane by mixing cell suspensions with petroleum ether for a short period (two minutes) (20). The quantification of mycolic acids through analysis of mycolic acid methyl esters showed levels similar to the wild type, concluding that the deletion of *Ag85A* shifts the TMM/TDM ratio. In contrast, Touchette et al. analyzed total cell mycolic acids and found higher accumulation of TMM and TDM in the deletion mutant compared to the wild type, and lower TDM synthesis using TMM with an N-alkyne moiety for click chemistry (40).

To quantify total TMM and TDM in the cell, we adopted Touchette et al.’s method to extract total membrane lipids from mycobacterial cells and confirmed the accumulation of TMM and TDM in the deletion strain (Fig.3A). For analyzing the localization of mycolic acids, we converted TMM and TDM to mycolic acid methyl esters (MAME; Fig. 3B) after fractioning them into arabinogalactan-associated and free TMM and TDM by delipidating. The results suggested that mycolic acids accumulated in the free lipid fraction in Δ*Ag85A* comparing with wild-type and complement strain (Fig 3B and Supplementary figure 3A). Since this fraction includes precursors, accumulation sites of mycolic acids could be the plasma membrane or mycomembrane.

**Figure 3.**
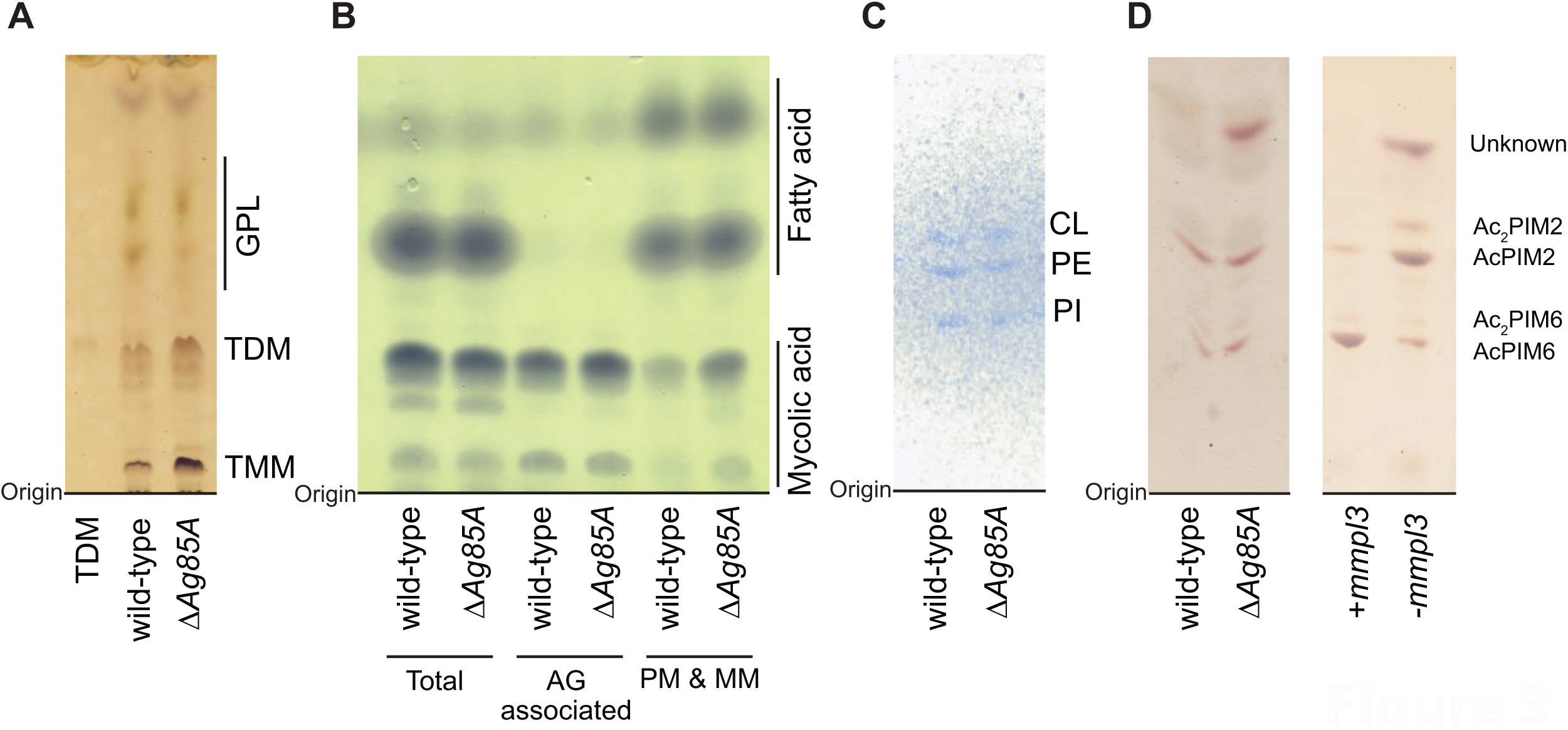
Mycolic acid and unknown glycolipid are accumulating in outer membrane of Δ*Ag85A.* (A) TMM and TDM were accumulated in Δ*Ag85A*. TMM and TDM were resuspended in chloroform, methanol, and water (9:1:0.1, v/v/v), chromatographed using chloroform, and visualized by orcinol. (B) Mycolic acid and fatty acid were methylated and chromatographed using a solvent containing petroleum ether and acetone (95:5, v/v). (C) Phospholipids were chromatographed using a solvent containing chloroform, methanol, 13 M ammonia, 1 M ammonium acetate, and water (180:140:9:9:23, v/v/v/v/v) and visualized by molybdenum blue (Rf = 0.35-0.61). (D) PIMs from the indicated mutants were separated as in the panel C (Rf = 0.20-0.48) and visualized by orcinol. *mmpl3* depletion strain was used for controls which accumulate mycomembrane precursor (TMM) in the plasma membrane. CL, cardiolipin; and PE, phosphatidylethanolamine. Pl, phosphatidylinositol. PIM, phosphatidylinositol mannoside. All experiments were repeated at least twice. Bands were assigned as the expected molecules based on migration patterns reported in previous literature.

Cell envelope precursors are synthesized in cytoplasm, anchored to inner leaflet of plasma membrane, flipped to outer leaflet, and transported to periplasmic region (41). If the deletion of *Ag85A* resulted in the accumulation of TMM in plasma membrane, such TMM accumulation may induce a stress response in the plasma membrane. To test this hypothesis, we analyzed both the phospholipids and the acylation status of phosphatidylinositol mannoside (PIM). Phospholipid levels are thought to play a role in sensing and responding to environmental stress affecting the plasma membrane (42). Additionally, we hypothesized that TMM accumulation may alter membrane fluidity, prompting *M. smegmatis* to modify or add acyl chains to PIM, as observed in response to membrane-fluidizing chemicals (43). Phospholipids including cardiolipin, phosphatidylethanolamine, and phosphatidylinositol were comparable in wild-type and Δ*Ag85A* (Fig. 3C). Acylation of PIM was also not altered in the deletion of *Ag85A* while the TMM accumulation by *mmpl3* knockdown caused the accumulation of di-acylated species of PIMs (Fig. 3D). Those data suggest that the deletion of *Ag85A* does not cause strong stress to the plasma membrane.

### Accumulated unsaturated and alkaline sensitive unknown glycolipids in ΔAg85A

In addition to TDM accumulation, TLC analysis revealed the presence of an unknown glycolipid band with a retention factor of 0.65–0.70 in the Δ*Ag85A* mutant, which was absent or present at low levels in the wild-type and complemented strains (Fig. 3D and Supplementary figure 3B). Glycolipids with high hydrophobicity are typically glycopeptidolipids (GPLs) or lipooligosaccharides (LOS), both components of the mycomembrane. Although LOS production has been reported to be absent in *Mycobacterium smegmatis* mc^2^155 strain (**44**), the genes required for LOS biosynthesis are conserved, and their expression may have been induced by the deletion of *Ag85A*. There is a possibility that could be influencing plasma membrane formation and membrane permeability. To characterize the unknown lipid, we performed alkaline hydrolysis to break down ester bonds in LOS and used iodine staining to visualize double bonds (Fig. 4B). The results indicated that the unknown lipid may contain unsaturated fatty acids and is sensitive to alkaline treatment. While these findings suggest that the unknown lipid could be LOS, further MALDI-TOF analysis did not show an increase in LOS signals (Supplementary figure 4), which appear as peaks at m/z 1689 and 1717 in LOS-producing *M. smegmatis* (**45**). Identification of this molecule remains to be determined.

**Figure 4.**
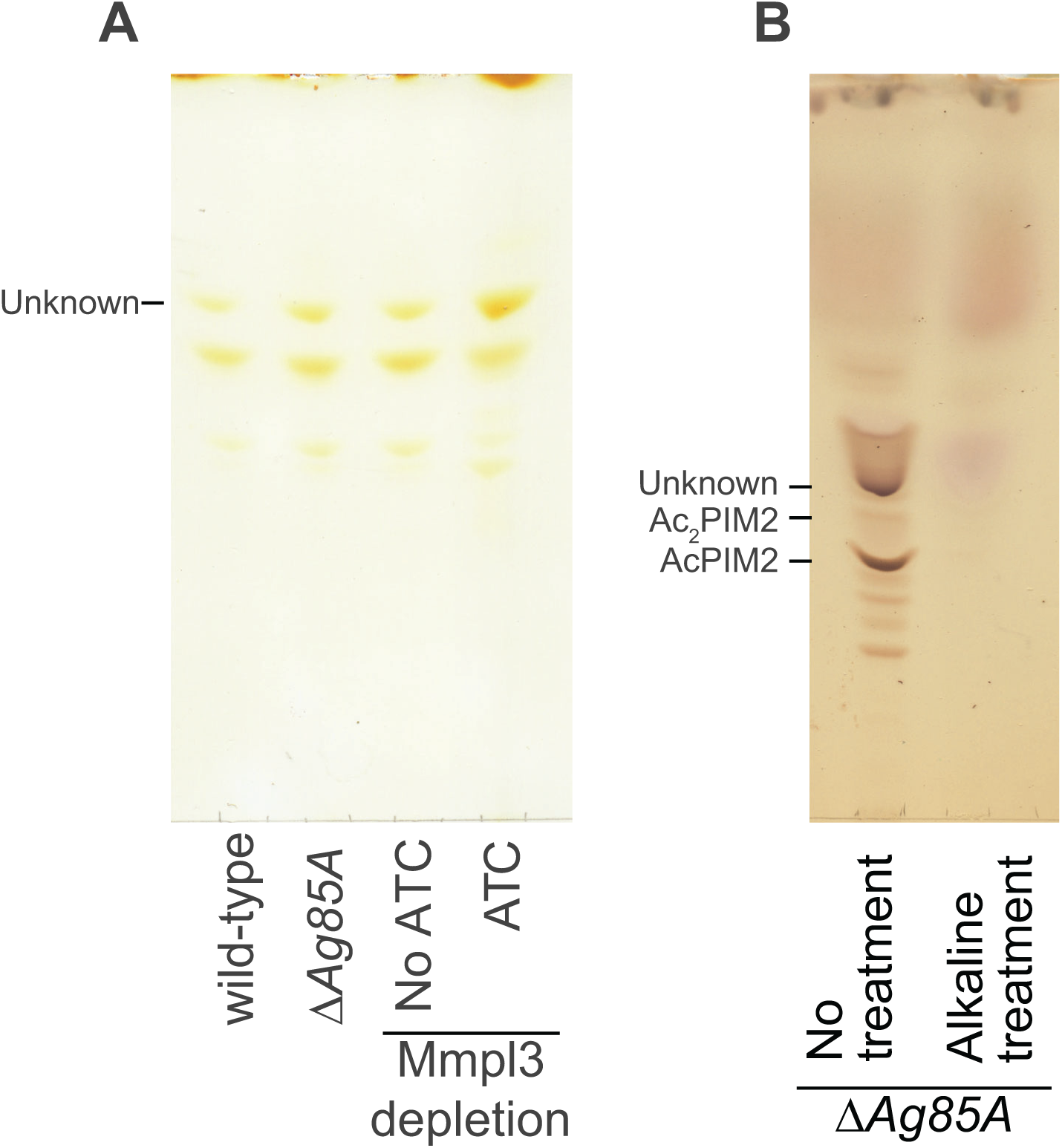
The unknown glycolipid is stained by iodine and susceptible to alkaline hydrolysis. (A) The unknown glycolipid was chromatographed using a solvent containing chloroform, methanol, 13 M ammonia, 1 M ammonium acetate, and water (180:140:9:9:23, v/v/v/v/v) and visualized by iodine vapor. (B) Lipid extracts were treated with 0.2M NaOH and resuspended in chloroform, methanol, and water (10:10:3, v/v/v). The plate was predeveloped by chloroform and developed by chloroform, methanol (9:1, v/v). Orcinol was used for visualization.

## Discussion

Ag85A is present across mycobacteria and plays a critical role in the synthesis of the mycobacterial cell envelope (20, 46). It is part of the antigen 85 complex, which is involved in the transfer of mycolic acids to the cell wall, essential for maintaining cell envelope integrity and pathogenicity (7–10, 47–49). Membrane domains are specialized regions in the plasma membrane that play a key role in coordinating the synthesis of cell wall peptidoglycan, certain phospholipids in plasma membrane, arabinogalactan, and outer mycomembrane lipids including phthiocerol dimycocerosate (27, 29, 50, 51). Feedback mechanisms from both the cell wall and phospholipids in the plasma membrane contribute to the regulation of plasma membrane domain formation, ensuring proper localization and function of these domains and thereby maintaining cell envelope integrity (29, 30, 52). Previous studies have demonstrated that the absence of *Ag85A* leads to hydrophilic micro-colony (20), increased sensitivity and accumulation of certain drugs (20, 21, 33), and changes in mycomembrane composition (40). In this study, we investigated the effects of *Ag85A* deletion using intrabacterial compound accumulation assays, membrane domain fractionation, fluorescence microscopy, and comprehensive lipid profiling. Our findings reveal that the absence of *Ag85A* disrupts membrane domain formation, likely due to the accumulation of free mycolic acids.

Based on previous findings and our current research, we propose that Ag85A has several key functions: 1) controlling TMM and TDM levels in the outer membrane, as the deletion led to the accumulation of these mycolates, primarily in the outer membrane (20, 40); 2) functioning within the feedback mechanism between accumulation of mycolic acids in outer membrane and membrane domain organization, as *Ag85A* depletion caused the dispersion of IMD marker proteins, which in turn disrupted cell envelope synthesis; and 3) facilitating the synthesis of other cell envelope components, since domain dispersion resulted in reduced cell envelope synthesis and increased membrane permeability. These data suggest that the outer membrane plays a role in the formation of plasma membrane domain in addition to the previously reported roles of the cell wall and plasma membrane phospholipids (30, 52).

Another possibility is that the accumulation of TMM/TDM increases the hydrophobicity of the outer mycomembrane, leading to disrupted molecular transport and diffusion. To compensate for this, the cell may produce the unknown glycolipid, which may function to balance membrane fluidity, possibly restoring molecular permeability in the mycomembrane. Such compositional changes may allow better diffusion and transport across the membrane despite the elevated hydrophobicity caused by TMM/TDM accumulation.

To further explore how *Ag85A* deletion leads to cell envelope disruption, we considered several mechanistic possibilities. One possibility is that TMM accumulates at the outer leaflet of the plasma membrane in Δ*Ag85A*. To investigate the site of free mycolic acid accumulation, we analyzed phospholipids and PIM acylation, using an *mmpl3* knockdown strain as a positive control. While *mmpl3* depletion induces PIM acylation, it is known to cause TMM accumulation specifically at the inner leaflet of the plasma membrane. However, it remains unclear whether TMM accumulation at the outer leaflet can similarly induce PIM acylation. Therefore, we cannot fully exclude the possibility that TMM accumulates at outer leaflet of plasma membrane in the absence of Ag85A. A second possibility is that the accumulation of free mycolic acids in Δ*Ag85A* results from compensatory activity by other Ag85 isoforms. Although knocking out *Ag85C* did not affect the protein levels of *Ag85A* and *Ag85B in M. tuberculosis* (53), it is still possible that *Ag85B* and/or *Ag85C* are upregulated in the absence of *Ag85A*. Such compensatory expression could contribute to the increased levels of free mycolic acids observed in the mutant.

A third possibility is that the deletion of Ag85A compromises the physical barrier function of the outer membrane. Previous studies have shown that Δ*Ag85A* forms less hydrophobic microcolonies (20) results in a permeable membrane (21), which may subject the plasma membrane to continuous stress from molecules that normally cannot cross the outer membrane. These observations support a model in which Ag85A contributes to maintaining the mycomembrane, modulating the expression of other *Ag85* isoforms, and sustaining the hydrophobic barrier function of the outer membrane—all of which may converge to influence membrane domain organization.

Maintaining the proper amount of mycolic acid in the mycomembrane is crucial for preserving membrane impermeability and stabilizing plasma membrane domains. By manipulating the mycolic acid in the mycomembrane, it may be possible to weaken the outer membrane, thereby increasing sensitivity to certain drugs, providing a potential strategy for targeting these organisms with antibiotics or other therapeutic agents. Plasma membrane domains were discovered decades ago, but their precise function and formation mechanisms remain speculative. Current and previous studies identified key genes involved in plasma membrane domain formation in *M. smegmatis*, including *ponA2* (a cell wall synthase), *cfa* (a C19-methylated fatty acid synthase), and *Ag85A* (an outer membrane synthase). Deletion mutants of these genes show significant disruptions in biofilm formation, impaired growth in macrophages *in vitro* and in mice *in vivo*, and increased drug sensitivity in pathogenic mycobacteria strains (19, 20, 22–25). These findings highlight the critical role of plasma membrane domains in maintaining cellular homeostasis and survival, and they also present plasma membrane domains as potential targets for new drug development.

## Acknowledgement

This work was supported in part by NIH grants R21AI144748 (to MSS and YSM), DP2AI138238 (to MSS), and U19AI142731 (to JSF), as well as by a fellowship from the Uehara Memorial Foundation (to TK). We thank the Mass Spectrometry Core Facility at the University of Massachusetts Institute for Applied Life Sciences, particularly Dr. Stephen Eyles, for technical support. We are also grateful to Dr. Kyoungtae Kim for sharing GraphPad Prism and to Dr. Babur Mirza for providing access to the bead beater.

## Material and Method

### Bacterial strains and growth conditions

Markerless knock-in *M. smegmatis* strains expressing both HA-mCherry-GlfT2 and Ppm1-mNeonGreen-cMyc or HA-mCherry-GlfT2 alone, and wild-type (mc^2^155) were previously established(27). Δ*Ag85A* mutant and the complemented strain wereere obtained from Dr. Anil Ojha (Wadsworth center, NY). Depletion strain of *Ag85A* was generated by introducing a CRISPRi construct provided from MSRdb(54) with electroporation(55). The cells were grown at 37°C in Middlebrook 7H9 broth supplemented with 11 mM glucose, 14.5 mM NaCl, and 0.05% (vol/vol) Tween 80. To deplete Ag85A, anhydrotetracycline (final concentration: 50 ng/mL; Cayman Chemical, Michigan, USA) was added and the culture was incubated overnight.

### Colony forming unit

Colony forming unit was analyzed as in the previous manuscript. In brief, Wildtype, Δ*Ag85A*, and complemented strains were grown to stationary phase, then diluted and grown to log phase. Cultures were treated with water (vehicle control) or 200 µg/mL dibucaine (MilliporeSigma, Massachusetts, USA) at 37°C for 3 hours, then cultures were serially diluted in 7H9 media and plated.

### CFU assay for ethambutol pretreatment

Cells were pretreated with either ethambutol or DMSO (vehicle control) for 3 hours. After incubation, the cells were washed to remove any residual ethambutol or DMSO. The cells were then treated with dibucaine or water for an additional 3 hours. CFU counts were obtained for each group both before and after dibucaine treatment. To compare the effects of the treatments, CFU ratios were calculated by dividing the CFU of the dibucaine-treated group by the CFU of the water-treated group. These ratios were determined both before and after treatment. Since CFU counts before treatment were expected to be within a similar range, the CFU ratio before treatment was normalized to 1.

### Intrabacterial drug/compound accumulation measurement

*M. smegmatis* (wild-type, Δ*Ag85A*, or the complement strain) cultures were grown in 7H9 broth supplemented with 11 mM glucose, 14.5 mM NaCl, and 0.05% (vol/vol) Tween 80, until to mid-log phase (OD_600_ ∼ 0.6). Cultures were then divided into three tubes for technical triplicate. Water (vehicle control) or dibucaine were added, and OD_600_ readings were taken at 0 hour, followed by incubation at 37°C with readings and sample collection at 1 hour. After incubation, OD_600_ was measured, and 1 mL samples were collected and pelleted by centrifugation at 13,000 rpm (14,549 rcf) for 10 min. Pellets were resuspended in 1 mL of 0.85% NaCl, pelleted again, and the supernatant was removed. For cell lysis, pellets were resuspended in 1 mL MAW solution Methanol/Acetonitrile/Water (2:2:1, vol/vol/vol) and stored at −80°C. Bead beating (6.5 m/s for 30s, 6 cycles, with ice cooling for 2 minutes between cycles) with Lysing Matrix B (MP Biomedical, California, USA) was performed to lyse cells (FastPrep24 5G, MP Biomedicals, OH), followed by centrifugation. Supernatants were transferred to 0.22 µm Spin-X centrifuge tube filters (Corning, New York, USA) and spun at 13,000 rpm (14,549 rcf) for 10–15 min at 4°C. Samples were stored at −80°C until LC-MS analysis.

### LC-MS analysis of intrabacterial accumulated drugs/compounds

The sample was analyzed using a Poroshell 120 EC-C18 column (2.1×50 mm; 1.9 µ particle size) and an Agilent 1290 Infinity II HPLC system connected to an Agilent InfinityLab LC/MSD system. The standard gradient program (shown as the percentage of acetonitrile) was used to build a solvent system with water/acetonitrile acidified with 0.1% formic acid: 0 to 0.4 min, 5%; 0.4 to 0.5 min, 5% – 30%; 0.5 to 3.0 min, 30% – 95%; 3.0 to 3.3 min, 95%; 3.3 to 3.6 min, 95% – 5%. At a flow rate of 0.5 mL/min and with a post-run flow of 1 min (post run is the period to allow the system to equilibrate for the next run at different time for 1.0).

Strong peaks for the analyte, or compound of interest, and e verapamil internal standard in this case, were verified in the calibration curve in standard mixtures of the compound in MAW before the biological samples were run. The lowest concentration of the analyte that could be identified and differentiated from the baseline so that the analyte peak height was at least twice the height of the baseline was known as the lower limit of detection (LLOD) for each compound. The linear-least squares method was used to determine the slope of the fitted line by plotting the compound/internal standard peak area ratio against the compound concentration in the calibration curve.

Another calibration curve was attained using the same method but substituting MAW with the cell lysate from an *M.* smegmatis culture grown in at least 15 mL media containing 7H9 broth supplemented with 10% ADS and 0.05% Tween 80 at 37 °C until reaching an OD_600_ ∼ 0.5 – 0.6. The bacteria were lysed by bead beating 6 cycles/2,400 rpm/30 s. The comparison of calibration curves with MAW vs. cell lysate was utilized to query the existence of a matrix effect which can be due to phenomena such ion suppression or enhancement^30^. Data processing and quantification of ions conducted were as published previously^14^. Briefly, for sample analysis, take 0.25 mL flow-through, followed by addition of verapamil, vortex, and inject 10 μL of the sample into the LC-MS. We extracted peak areas of dibucaine and verapamil in dibucaine and DMSO treated samples and calculate the drug/internal standard (D/I) ratio, plotting the calibration curve using linear regression (y = mx) and divide by the calibration factor. Normalize by OD_600_ at the time point and calculate accumulation for three replicates.

### Membrane fractionation

Membrane fractionation was proceeded as in the previous literatures In brief, 5 mL of seed culture was inoculated into three 2-L glass flasks, each containing 500 mL of supplemented 7H9 medium, and incubated at 37°C for 16–18 hours. At mid-log phase (OD₆₀₀ = 0.5–1.0), the culture was transferred to 1,000-mL bottles and centrifuged at 8,000 rpm for 15 minutes at 4°C using an F9S-4×1000Y rotor. The resulting pellet was resuspended in 50 mM HEPES/NaOH buffer (pH 7.4) and transferred to pre-weighed 50 mL conical tubes. The samples were then centrifuged at 4,000 rpm for 10 minutes at 4°C using an Eppendorf A-4-81 rotor. After centrifugation, the supernatant was removed, and the pellet was resuspended in 40 mL of 50 mM HEPES/NaOH buffer (pH 7.4). The samples were centrifuged again under the same conditions, and the final pellet was weighed.

The pellet was then resuspended at a ratio of 1 g per 5 mL in lysis buffer containing 1/25 volume of protease inhibitor mix (Roche). Cell disruption was carried out by nitrogen cavitation at 1,850–2,200 psi for 30 minutes, with pressure released every 10 minutes, repeated for a total of three cycles. A sucrose gradient (20–50% w/v) was prepared using a BioComp Gradient Master. Following cavitation, the lysate was centrifuged at 4,000 rpm for 10 minutes at 4°C, and the supernatant was carefully transferred to a new tube, avoiding the top layer. This centrifugation and transfer step was repeated once. A volume of 1.2 mL of clarified lysate was carefully layered onto the sucrose gradient and centrifuged at 35,000 rpm (218,000 × g) for 6 hours at 4°C using a Beckman L8-70M ultracentrifuge equipped with an SW40 swing rotor. Gradient fractions were collected using a BioComp gradient collector, and 450 µL from each fraction was aliquoted into separate tubes, and stored at –80°C.

### SDS-PAGE and western blotting

SDS-PAGE and western blotting were proceeded as in the previous literatures(26). For sample preparation, 12 µL of each fraction was mixed with 4 µL of 4× Sample Loading Buffer and incubated at 95°C for 5 minutes. Protein ladder was prepared by mixing 10 µL of Precision Plus Protein Kaleidoscope Standards (Bio-Rad, California, USA) in an Eppendorf tube and incubating at 95°C for 5 minutes. The samples and ladder were then loaded onto SDS-PAGE gels and run at a constant current of 25 mA per gel.

Following electrophoresis, proteins were transferred onto membranes at 14 V overnight using a Bio-Rad PowerPac 1000 system (Bio-Rad, California, USA). Membranes were blocked in 5% skim milk prepared in PBS containing 0.05% Tween 20 (PBS-T) for 1 hour at room temperature. Blots were incubated overnight at 4°C with rabbit polyclonal anti-PimB’ or anti-MptC antibodies (56). The membranes were then incubated with horseradish peroxidase-conjugated secondary antibodies—either donkey anti-rabbit (Cytiva) or sheep anti-mouse (Millipore Sigma, California, USA)—at a 1:4,000 dilution. Protein bands were visualized using a homemade enhanced chemiluminescence (ECL) reagent, and images were captured using the Amersham ImageQuant 800 system (Cytiva).

### Microscopy and image analysis

Microscopy and image analysis were performed in the previous literatures(30). In brief, a 5 µL aliquot of bacterial culture was placed on an agar pad (1% agarose in water) positioned on a glass slide. Images were captured using a Nikon Eclipse E600 microscope (Nikon Eclipse Ti with ×100 objectives, N.A.=1.30). Cell outlines were traced using the Oufti software(58). MATLAB scripts were used to generate intensity profiles(50). Cap-to-side wall ratios were determined by summing signal values from the 15% of the cell length at each pole and comparing this by the signal values from the 70% of mid-cell region.

### Lipid extraction and analysis

Analysis of MAME was performed as described before (59). Briefly, 50 mg of mycobacterial cells are mixed with 2 mL of 15% tetrabutyl ammonium hydroxide and heated overnight at 100°C. The next morning, the mixture is diluted with 2 mL water, 1 mL dichloromethane, and 250 μL iodomethane, then stirred for 30 minutes. The upper aqueous layer is discarded, and the lower organic layer is washed with 3 mL of 1 M HCl followed by 3 mL of water. If needed, the volume is reduced. The residue is dried under a stream of nitrogen gas, dissolved in dichloromethane, transferred to a polypropylene tube, and evaporated under nitrogen. The dried MAMEs are dissolved in 0.2 mL toluene and 0.3 mL acetonitrile, incubated at 4°C for 1 hour, and any insoluble debris is discarded to yield the MAME extract. For TLC, the MAMEs are spotted on aluminum-backed silica sheets, developed in petroleum ether/acetone (95:5, vol/vol), and visualized by spraying with 5% ethanolic phosphomolybdic acid followed by charring at 110°C. Arabinogalactan associated or non-associated mycolic acid were separated by dilapidation in chloroform/methanol (2:1, vol/vol).

Analysis of phospholipids and glycolipids were performed in the previous literature. In brief, 50 OD_600_ units of cells was harvested into a pre-weighed 50-mL conical tube, centrifuge at 4000 rpm for 10 minutes, discard the supernatant, and repeat to collect all cells into one tube. Remove as much supernatant as possible using an aspirator, then weigh the tube to calculate the pellet weight. Add 20 volumes of chloroform/methanol (2:1) and extract for 1.5 hours at room temperature, ensuring the pellet is fully suspended. Spin and transfer the supernatant to a fresh 15-mL conical tube. Repeat the extraction with 10 volumes of chloroform/methanol (2:1), spin, and transfer the supernatant again. Perform a third extraction with chloroform/methanol/water (1:2:0.8), spin, and combine the supernatants. Dry 800 µL of the combined organic solvent under nitrogen, resuspend in 100 µL water, and add 200 µL of water-saturated butanol. Vortex, centrifuge for 30 seconds, and transfer the upper phase to a new tube. Repeat the butanol extraction, then add 100 µL water to the combined butanol phase and repeat. Transfer the upper phase to a new tube, dry it using a speed-vac, and resuspend in 20 µL of water-saturated butanol. For TLC, equilibrate the tank with chloroform/methanol/13 M NH_3_/1 M NH_4_Ac/water (180:140:9:9:23), spot 3 µL of sample on an HPTLC plate (MilliporeSigma, California, USA), and develop for ∼45 minutes. Dry the plate and stain using orcinol reagent at 100°C for 5 minutes, cupric acetate at 150°C for 10-20 minutes, iodine vapor for 10-20 minutes at room temperature, or molybdenum blue reagent for 10 minutes at room temperature.

### Alkaline hydrolysis of lipid extract

Dry the desired volume of lipid-containing solution under a nitrogen stream, then resuspend it in 100 µL of 0.2 M NaOH in methanol or methanol (methanol only treatment) and incubate at 40°C for 2 hours. Neutralize the reaction with 1.2 mL of glacial acetic acid and dry under nitrogen. Resuspend the dried sample in 100 µL of water, add 200 µL of water-saturated butanol, vortex, centrifuge for 30 seconds, and transfer the upper phase to a new tube. Repeat the extraction with an additional 100 µL of water-saturated butanol. Add 100 µL of water to the combined butanol phase, repeat the extraction, and dry the final butanol phase using a speed-vac. Resuspend the dried sample in 10 µL of chloroform/methanol/water (10:10:3, vol/vol/vol), wash the tube with 5 µL of chloroform/methanol (2:1, vol/vol), and load onto a TLC plate. Pre-develop the TLC plate with 100% chloroform, then develop it with chloroform/methanol (9:1, vol/vol). Once dry, spray the plate with orcinol reagent, bake at 100°C for 5 minutes, and save the image using a computer scanner.

### MALDI-TOF-MS Analysis of lipid extract

Lipid extracts were analyzed by matrix-assisted laser desorption/ionization time-of-flight mass spectrometry (MALDI-TOF-MS). The 1-butanol extract was mixed in a 1:1 volume ratio with a saturated solution of 2,5-dihydroxybenzoic acid (DHB) prepared in 70% acetonitrile with 0.1% trifluoroacetic acid. Samples were spotted onto a stainless steel MALDI target plate and allowed to air-dry at room temperature. Mass spectra were acquired using a Bruker UltrafleXtreme MALDI-TOF/TOF mass spectrometer (Bruker Daltonics, Billerica, MA, USA) operated in linear positive-ion mode, with an accelerating voltage of 20,000 V. Spectra were collected by averaging 500 laser shots per sample. External calibration was performed using a peptide or lipid standard mixture to ensure mass accuracy.

